# Brief report: Mutations in SIV Nef that disrupt and restore tetherin downregulation

**DOI:** 10.1101/833137

**Authors:** Nicholas J Maness, Blake Schouest

## Abstract

The H_196_ residue in SIVmac239 Nef is conserved across nearly all HIV and SIV isolates, lies immediately adjacent to the AP-2 (adaptor protein 2) interacting domain (ExxxLM_195_), and is critical for several described AP-2 dependent Nef functions, including the downregulation of tetherin (BST-2/CD317). Surprisingly, many stocks of the closely related SIVmac251 swarm virus harbor a *nef* allele encoding a Q_196_, which is associated with loss of multiple AP-2 dependent functions in SIVmac239. Publicly available sequences for SIVmac251 stocks were mined for variants linked to Q_196_ that might compensate for functional defects associated with this mutation. Variants were engineered into the SIVmac239 parental plasmid and mutant viruses were used to test tetherin downregulatory capacity in primary CD4 T cells using flow cytometry. SIVmac251 stocks that encode a Q_196_ residue in Nef uniformly also encode an upstream R_191_ residue. We show that R_191_ restores the ability of Nef to downregulate tetherin in the presence of Q_196_. However, a published report showed Q_196_ commonly evolves to H_196_ in vivo, suggesting a fitness cost. R_191_ may represent compensatory evolution to restore the ability to downregulate tetherin lost in viruses harboring Q_196_.

## Introduction

The lentiviral Nef protein is a common target of CD8-T lymphocyte (CD8TL) responses in both HIV-1 infected persons and SIV infected rhesus macaques and readily evolves to evade these responses [1–6]. Nef is highly pleiotropic and mediates the downregulation of several cell surface molecules involved in innate and adaptive immune responses against virus infected cells such as TCR-CD3 (in most SIVs but not HIV-1) [7], CD4 [8–10], CD8αβ [11], CD28 [12], tetherin (BST2 or CD317; in most SIVs and in HIV-1 group O, but not HIV-1 group M) [13–15], MHC-I [16], MHC-II [17], CD1d [18], CD80/CD86 [19] and likely others as well as enhancing viral infectivity by preventing virion incorporation of host serine incorporator 3 (SERINC3) and SERINC5 proteins [20–23]. Nef-mediated modulation of several of these molecules, including CD4, CD8αβ, CD28, tetherin, and SERINC3 and SERINC5 requires interactions between Nef and adaptor protein (AP)-2 complexes [11, 20, 24–28].

We used high throughput next generation sequencing to track evolution in SIV Nef [29, 30], with particular focus on viral escape from antiviral CD8TL responses, including CD8TL targeting the SIV Nef IW9 (IRYPKTFGW_173_, with subscript numbers representing the position in the SIVmac239 Nef protein) and MW9 (MHPAQTSQW_203_, hereafter referred to as MW9) epitopes in rhesus macaques that express Mamu-B*017:01. MW9 overlaps the well-defined “di-leucine” ExxxLM_195_ motif and lies immediately upstream of the DD_205_ di-acidic motif also important for AP-2 binding [31]. Though selection eventually favored changes of the first position in MW9, specifically M_195_I or M_195_V, an H_196_Q (second position in MW9) substitution was initially favored in several animals. Since this variant was never fixed and generally lost soon after arising, we hypothesized it may have represented an effective escape mutation yet imparted a negative impact on Nef function. Specifically, we tested whether functions involving interactions with AP-2 would most likely be impacted, given the close proximity of this epitope with the ExxxLM_195_ AP-2 interaction domain. Not surprisingly, the H_196_Q variant selectively disrupted Nef functions that rely on interactions with AP-2, such as downregulation of tetherin, CD4, and CD28 and disrupted Nef’s ability to reduce SERINC5-mediated reductions of viral infectivity, while having no impact on MHC-I or CD3 downregulation, functions that do not rely on AP-2 interactions [32, 33]. In that study, we did not identify any potential compensatory mutations that allowed for regain of function in the presence of the H_196_Q variant leading to this variant being only fleetingly detected and eventually replaced by escape mutations with less significant impacts on important Nef functions.

Many isolates of SIVmac251, a commonly used strain in SIV studies, harbor a Q_196_ in the viral Nef protein. In this study, we sought mutations linked to Q_196_ that might compensate for loss of function associated with this residue. We then used publicly available viral sequences to determine if Q_196_ was stable in vivo. We identified an upstream variant, R_191_ (E_191_ in SIVmac239) that compensates for the loss of tetherin downregulation associated with Q_196_. However, we also found that Q_196_ routinely mutated to H_196_ in vivo, suggesting reduced fitness despite the maintenance of tetherin downregulation associated with the combination of Q_196_ and R_191_ residues.

## Materials and Methods

### Ethics Statement

Cells used in this study were taken from blood from six Indian-origin rhesus macaques (*Macaca mulatta*) that are part of the breeding colony at the Tulane National Primate Research Center. Animals were anesthetized as part of their routine semi-annual health assessment (SAHA) and additional blood was drawn for this study. Thus, animals were not anesthetized specifically for the studies described herein. All animals were housed in compliance with the NRC Guide for the Care and Use of Laboratory Animals and the Animal Welfare Act. Blood draws were approved by the Institutional Animal Care and Use Committee of Tulane University (OLAW assurance #A4499-01) under protocol P0191. The Tulane National Primate Research Center (TNPRC) is fully accredited by AAALAC International [Association for the Assessment and Accreditation of Laboratory Animal Care(AAALAC#000594)], Animal Welfare Assurance No. A3180-01. Breeding colony animals at the TNPRC are housed outdoors in social groups and frequently monitored by veterinarians and behavioral scientists. The animals were fed commercially prepared monkey chow and supplemental foods were provided in the form of fruit, vegetables, and foraging treats as part of the TNPRC environmental enrichment program. Water was available at all times through an automatic watering system. The TNPRC environmental enrichment program is reviewed and approved by the IACUC semiannually. Veterinarians at the TNPRC Division of Veterinary Medicine have established procedures to minimize pain and distress through several means. Monkeys were anesthetized with ketamine-HCl (10 mg/kg) or tiletamine/zolazepam (6 mg/kg) prior to all procedures. The above listed anesthetics were used in accordance with the recommendations of the Weatherall Report.

### Primary cell isolation, culture and infection

Primary CD4 T cells were magnetically isolated from PBMC from healthy rhesus macaques using nonhuman primate CD4 microbeads (Miltenyi) according to the manufacturer’s protocol. Isolated cells were stimulated with concanavalin A for two days and cultured thereafter with R15/50 media, comprised of RPMI media with 15% FBS and 50U/ml IL-2. SIV infections were conducted using the spinoculation technique [34] with each 1ml aliquot of virus (approximately 10^8 viral RNA copies per milliliter) layered on 100ul of 20% sucrose solution and centrifuged for 1 hour at 4°C at 20,000xg. After removal of the supernatant, the concentrated virus was resuspended in 100ul of R15/50 media and gently dripped onto one million CD4 T cells plated at 1 million cells/ml in 48 well plates. Plates were then spun at 2000rpm for 2 hours at room temperature. After centrifugation, plates were placed in 37C humidified incubators with 5% CO_2_. Cells were cultured for 36 hours before harvest for flow cytometry assays.

### Mutant virus production

Mutants of the SIVmac239 virus were generated using site directed mutagenesis of the SIVmac239 3’ hemiplasmid, using mutagenesis primers designed using web-based software (PrimerX from bioinformatics.org). To generate the H_196_Q mutation alone, we used the following mutagenesis primers using the QuikChange II Site-Directed Mutagenesis Kit (Agilent); F: GCA TTA TTT AAT GCA GCC AGC TCA AAC TTC CC, and R: GGG AAG TTT GAG CTG GCT GCA TTA AAT AAT GC and to generate the E_191_R mutation on the backbone that already contained the H_196_Q variant, we used the following mutagenesis primers; F: GGC ACA GGA GGA TGA GAG GCA TTA TTT AAT GCA GC, and R: GCT GCA TTA AAT AAT GCC TCT CAT CCT CCT GTG CC. After mutagenesis, plasmids were treated with DpnI to remove non-mutated parental plasmids and cloned into Stbl2 cells (Life Technologies) using the manufacturer’s protocol. Mutations were sequence-verified, and successfully mutated plasmids were used for follow-up studies. Mutated 3’ plasmids were ligated with the 5’ hemiplasmid, transfected into Vero cells followed by harvest of the virus-containing supernatant. Of note, the Nef region of interest overlaps with the 3’ long terminal repeat (LTR). Attempts to mutate the full-length SIV plasmid resulted in mutations in both the 5’ and 3’ LTR regions, which rendered the viruses replication incompetent, necessitating the need for mutating only the 3’ hemiplasmid followed by ligation to the wild type 5’ plasmid. Some viruses were further expanded in CEMx174 cells. Viral RNA was extracted from all viral stocks and sequenced to ensure the presence of desired mutations.

### Flow cytometry

Thirty-six hours after infection, primary cells were stained with labeled antibodies to CD4 (BV421, clone L200, BD Biosciences) and tetherin (PE, clone RS38E, Biolegend), followed by fixation, permeabilization, and intracellular labeling with a FITC labeled antibody against the Gag p27 protein (clone 55-2F12). Data was acquired on a BD LSRII instrument and analyzed using Flowjo v10 software.

### Structural analysis

Structures showing interactions between SIVsm Nef, Tetherin, and AP-2 subunits were recently published [35]. We used UCSF Chimera software [36] to probe potential interactions between our Nef residues of interest at positions 191 and 196 and host AP-2 and tetherin proteins. The Rotamers function in UCSF Chimera was used to predict impacts of mutations and the Matchmaker function was used to assess positioning of Nef amino acids in SIV relative to HIV-1.

### Sequence analysis and alignments

Nef sequences from a broad array of SIV isolates were identified from a published report [37] and downloaded from NCBI for amino acid alignments using Geneious Prime 2019.1.3 using the built-in Geneious Alignment algorithm with default settings. SIVmac251 sequences available from published reports [38, 39] were downloaded from NCBI into Geneious Prime 2019.1.3 and mapped to SIVmac239, used as the reference genome, followed by identification and quantification of variations using the Find Variations/SNPs function. Sequences published in the Lamers et al report [39], were first divided into those extracted from the inoculum and from individual tissues, which were analyzed separately.

## Results

### Conservation of the Nef H_196_ residue among primate lentiviruses

To assess conservation of the H_196_ residue, we performed alignments of the flexible loop region of Nef from a diverse array of SIV isolates from Africa. Although the flexible loop is, in general, far more variable than the core, we found the H_196_ residue to be almost completely conserved among all isolates sequenced to date (Figure 1A), even more conserved than important residues in the adjacent “di-leucine” ExxxLM_195_ motif, E_190_ and L_194_.

**Figure 1.**
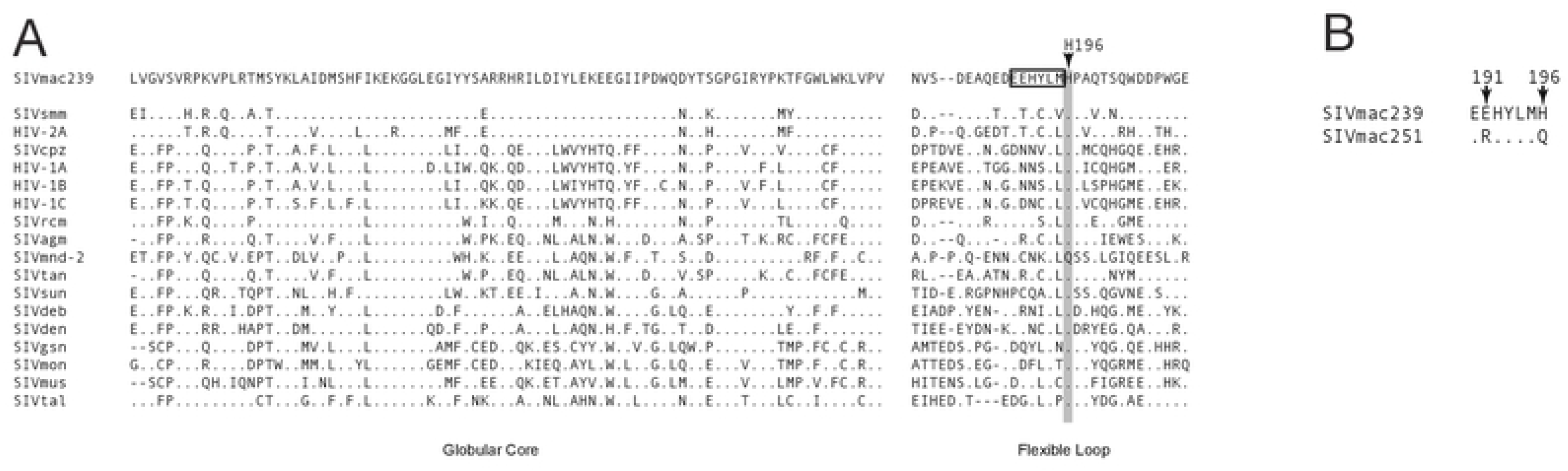
Assessment of the conservation of residue 196 in SIV Nef. Alignment of the Nef core and flexible loop from several strains of SIV (A). Sequences were derived from a recent study [37]. The ExxxLM AP-2 binding motif is boxed and the H_196_ residue (based on SIVmac239 numbering) is highlighted and noted by the arrow. SIVmac251 stock sequences from a published study [38] show a variant amino acid upstream of Q_196_ (R_191_) that was always associated with Q_196_ (B). Alignments were performed using Geneious Prime 2019.1.3.

We next scanned publicly available sequences from a recent study that used single genome amplification to extensively examine SIVmac251 challenge stocks [38]. In this report, a Q_196_ residue was detected in a large fraction of sequences from SIVmac251 stock viruses from several labs. Interestingly, there was a perfect linkage between the Q_196_ residue and an upstream R_191_ residue, which is E_191_ in SIVmac239 (Figure 1B). Of 38 total sequences that contained the region of interest, derived from three different challenge stocks, 25 sequences contained both R_191_ and Q_196_ while Q_196_ was never found in the absence of R_191_. Other nearby variants relative to SIVmac239 were detected but only R_191_ co-occurred with Q_196_ in all sequences.

### Upstream compensatory variant restores tetherin downregulation

Given the strong linkage between the R_191_ (E_191_ in SIVmac239) variant and Q_196_, we tested whether R_191_ allowed tetherin downregulation in the presence of Q_196_. The R_191_ residue lies within the ExxxLM_195_ motif (EEHYLM_195_ in SIVmac239, E<u>R</u>HYLM_195_ in many SIVmac251 isolates). We introduced the E_191_R variant along with the H_196_Q onto the SIVmac239 backbone to assess tetherin downregulation. Viruses harboring H_196_Q alone were largely deficient in tetherin downregulation, as expected. When E_191_R was introduced along with H_196_Q, the resulting virus showed full competency in tetherin downregulation, similar to SIVmac239 (Figure 2A). N-fold analysis of tetherin downregulation in cells from multiple animals demonstrated significant loss of downregulation in the virus harboring only H_196_Q, while the addition of E_191_R restored this ability to wild type levels (Figure 2B).

**Figure 2.**
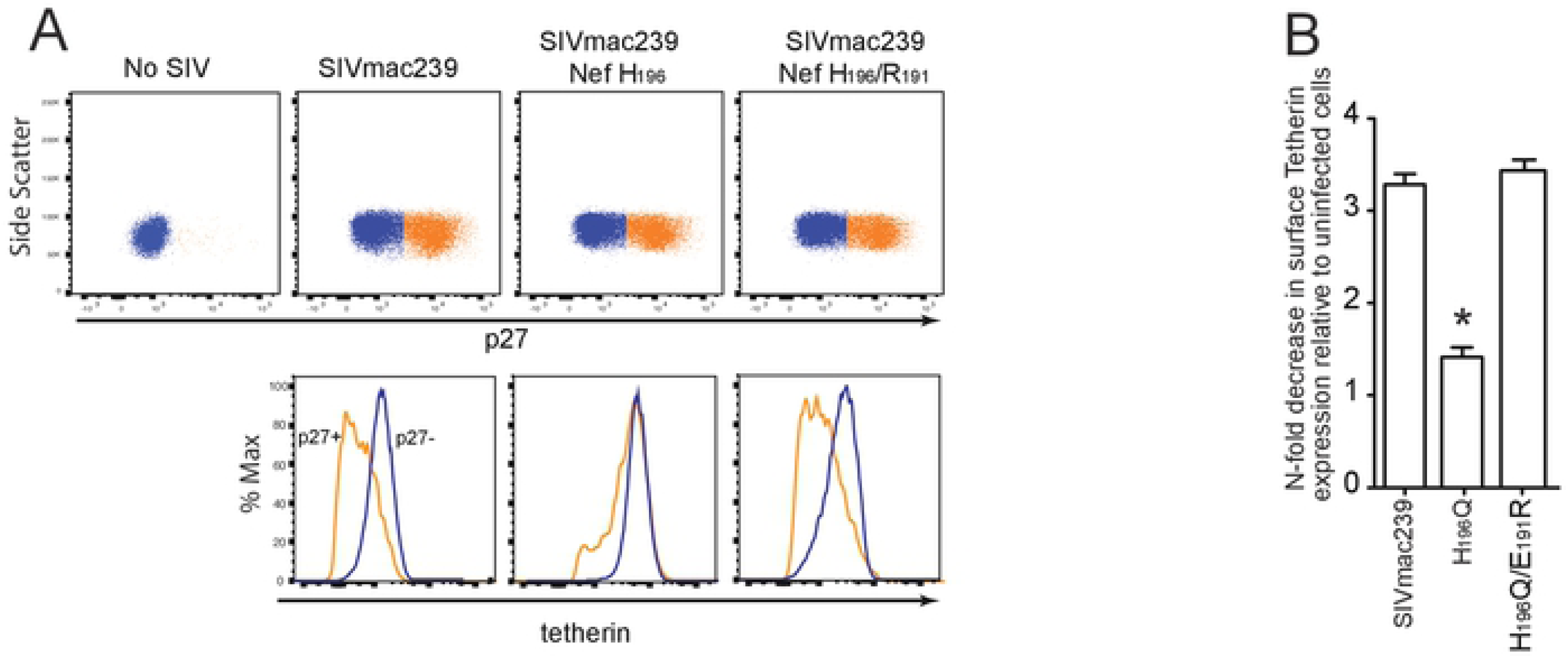
SIVmac251 maintains tetherin downregulation despite a unique AP-2 binding motif. (a) Representative flow cytometric analysis of surface expression of tetherin on primary CD4 T cells infected with wild type SIVmac239 or SIVmac239 harboring the H_196_Q variant alone or in combination with the E_191_R variant. Cells were identified as infected via intracellular Gag p27 staining, top row, as we have described previously [29, 30]. Infected cells (p27+) are orange, while uninfected (p27−) are blue. Surface expression of tetherin compared between infected cells (orange line) and uninfected (blue line) are shown in the bottom panels. (b) N-fold analysis of tetherin downregulation from multiple experiments using cells derived from at least three different RM and compared by way of a two-tailed t-test.

### Structural insights

The structure of SIVsm Nef bound to simian AP-2 was recently published [35]. We used UCSF Chimera structural analysis software [36] to assess how the residues at positions 191 and 196 interact with host AP-2 and tetherin molecules. The critical residues in the dileucine motif [E_190_, L_194_, V_195_ (M_195_ in SIVmac239)] show clear interaction with AP-2, while H_196_ is oriented in the opposite direction (Figure 3A), similar to H_166_ in HIV-1 (Figure 3B) [28], which is homologous to H_196_ in SIV. We next used the Rotamers function in UCSF Chimera to determine whether the H_196_Q variant impacted interactions with AP-2. Replacement of the H with a Q at this position resulted in a large number of possible rotamers for Q_191_, nearly all of which maintained a similar orientation as H_191_, directed away from AP-2, suggesting no obvious impact on the interaction between Nef and AP-2. However, the H_196_Q variant is predicted to disrupt a salt bridge between H_196_ and tetherin residue D_15_ as assessed using PISA (Proteins, Interfaces, Structures, and Assemblies) software [40], suggesting disruption of a direct interaction between Nef and tetherin may contribute to the selective disadvantage of this change.

**Figure 3.**
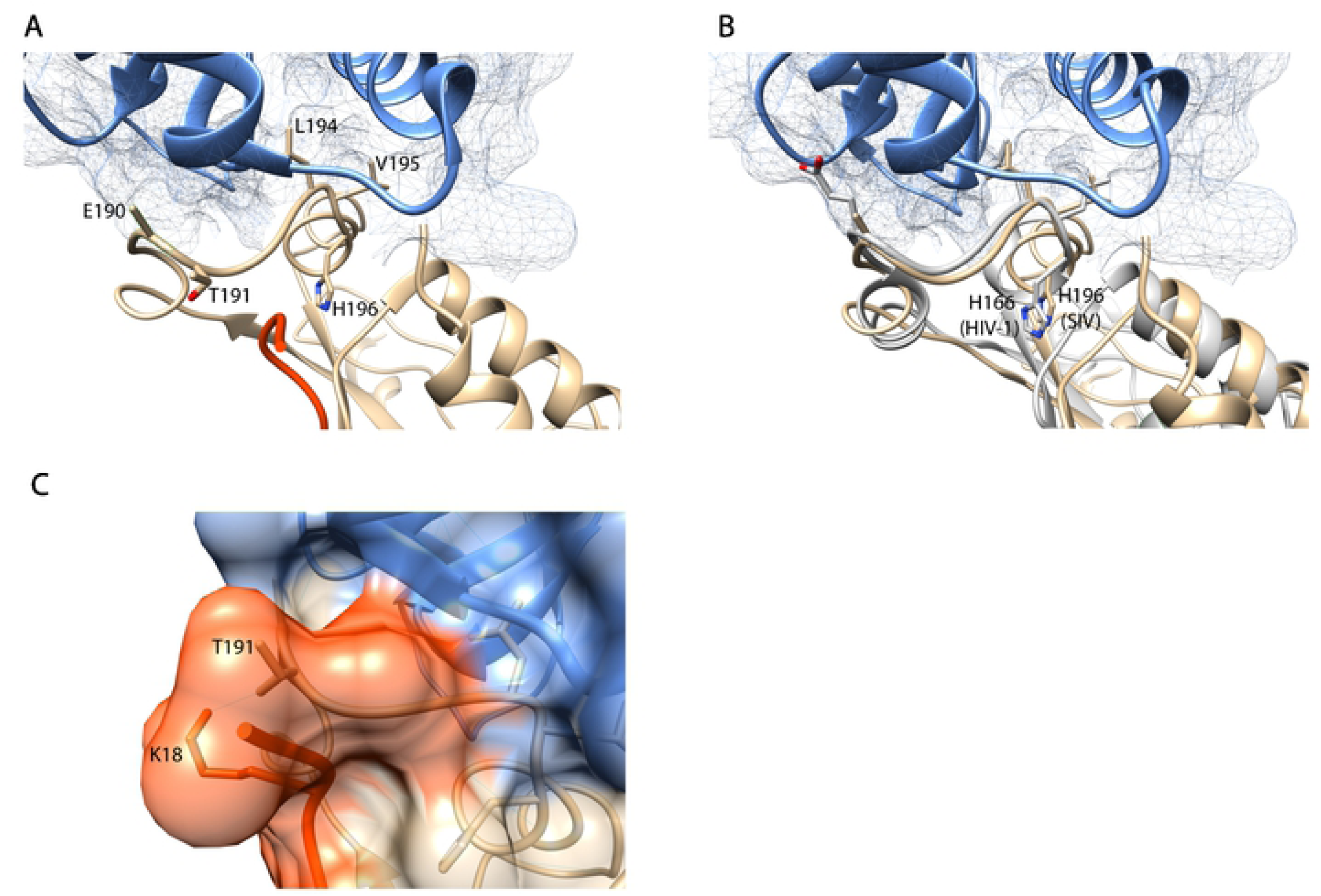
Structural insights into Nef, AP-2, and tetherin interactions. (a) Although adjacent to the ExxxLM motif that directly binds Nef to AP-2, H196 is oriented away from this interaction. (b) Alignment between SIVsm and HIV-1 Nef (PDB: 4NEE) with H_166_ (HIV-1) and H_196_ (SIVsm) highlighted. (c) T_191_ in SIVsm directly interacts with K_18_ in the DIWK motif of tetherin.

Position 191 is a T in SIVsm, as opposed to E_191_ in SIVmac239. While this residue does not contact AP-2, intriguingly, it does interact directly with the K_18_ residue in the DIWK motif of the tetherin protein itself via a hydrogen bond (Figure 3C). Replacement of T_191_ with an E (as in SIVmac239) maintained the predicted hydrogen bond with K_18_, suggesting this interaction holds true between SIVmac239 and tetherin. Further, replacement of E_191_ with an R resulted in only low probability orientations, preventing a meaningful analysis of this structural change.

### In vivo stability of the Q_196_ residue

Finally, we wished to assess the in vivo stability of the Q_196_ residue. We hypothesized that since the combination of Q_196_ and R_191_ residues allowed for efficient tetherin downregulation in vitro, that Q_196_ would be stable in vivo when it exists in combination with R_191_. We used publicly available sequences from a recent study wherein macaques were infected with an stock of SIVmac251 that harbored virus with nearly 90% containing the combination of Q_196_ and R_191_ [39] based on our analysis of their deposited sequences. That study used a modified Single Genome Amplification (SGA) method to quantify viral variation in plasma throughout infection and multiple neurological tissues at necropsy. After infection, the Q_196_ residue was detectable primarily at three weeks post infection, with the exception of a small number of reads that contained Q_196_ at 3 months. This residue was thereafter lost in all three animals and was not detected in any neurological sites in any animals at necropsy (meninges, parietal lobe, temporal cortex) (data available in the cited manuscript and in their deposited sequences).

## Discussion

The SIVmac251 viral swarm is pathogenic in rhesus macaques and has been used in hundreds of studies to date. However, this swarm has been independently grown in many labs using multiple cell types and under a variety of conditions [38]. It stands to reason that there may be genetic differences between SIVmac251 viral stocks leading to unique biological differences, but few of these differences have been characterized for how they impact specific virologic properties, including the downregulation of host tetherin.

The ability to downregulate host tetherin is a feature of a wide variety of enveloped viruses ranging from Ebola to HIV [41, 42]. Most SIV isolates use the viral Nef protein to perform this task [13, 43, 44] but several isolates use alternate pathways, suggesting strong selection to maintain this function. Surprisingly, Nef encoded by SIVcpz cannot downregulate human tetherin and studies suggest that evolution of HIV-1 Vpu to gain the ability to downregulate tetherin was a critical event in the HIV-1 epidemic [43, 45]. Thus, countering tetherin likely is an important feature of all or nearly all SIV and HIV isolates.

In addition to interactions with host AP-2 proteins, SIV Nef is known to interact directly with the tetherin protein [46] and a subset of those interactions were recently verified structurally [35]. These structures show that H_196_ does not directly interact with host AP-2 but is predicted to form a salt bridge with tetherin, which is predicted to be disrupted in the H_196_Q variant using PISA software [40]. However, our previous report showed that the H_196_Q variant disrupted multiple Nef functions that rely on AP-2 interactions suggesting that disruption of a direct interaction with tetherin likely does not fully explain the functional deficits identified in this variant. Here, we show that evolution of the E_191_R variant restored tetherin downregulation in the presence of Q_196_. Intriguingly, T_191_ in SIVsm interacts directly with the lysine in the DIWK motif in the tetherin protein [35], suggesting variation at this residue may impact tetherin downregulation via a direct effect on this interaction. E_191_ in SIVmac may also interact with this K_18_ residue as these two amino acids are well known to form hydrogen bonds, although we cannot confirm without structural data.

Many strains of SIVmac251 encode a Nef protein with a Q_196_ residue, which is always linked to an upstream R_191_ residue. Here we show that the presence of R_191_ fully restores competency in downregulation of tetherin in the presence of Q_196_. Nonetheless, our data also suggest that Q_196_ is not stable in vivo and evolves to H_196_, the residue present in nearly all SIV isolates. These data beg the question of how the Q_196_ residue arose in the first place. It’s possible it arose during replication in cultured cells where selection pressures are undoubtedly different than those the virus experiences in vivo. Given our data suggesting the H_196_Q variant can arise in vivo in SIVmac239 infected macaques as a result of escape from CD8TL responses [30], these data may suggest that SIVmac251 was isolated from an animal that targeted this region with CD8TL, leading to viral escape, and prior to other escape variants becoming dominant, as happened in our previous study [30].

Taken together, our mutational and functional data combined with published structural and sequence data suggest the possibility that the E_191_R variant in SIVmac might enhance an interaction between Nef and tetherin thus restoring the ability of Nef to downregulate tetherin in the presence of the H_196_Q variant, but that this variant may not restore other functions, thus leading to selection to restore H_196_ in vivo. The existence of compensatory variation in viral proteins has been described in SIV and HIV-1 [47–50] but those descriptions are restricted to viral structural proteins, primarily Gag. Our data suggest R_191_ in SIVmac251 may exist to compensate for loss of function associated with Q_196_. If so, this may be the first report of compensatory variation in the viral Nef protein or any nonstructural viral protein. However, our analysis of published in vivo data clearly demonstrate that Q_196_ evolves to H_196_ in vivo. Finally, our data do not suggest that stocks of SIVmac251 that harbor a Q_196_ residue are in any way less useful than stocks that do not. Instead, our data underscore the need to understand the evolutionary pressures that give rise to particular viral variants, which may be relevant in the choice of virus stock for animal model experiments.

## Author contributions

NJM conceived the study and wrote the first draft. BS conducted the experiments and edited all drafts.

